# Nano-CT characterization reveals coordinated growth of a rudimentary organ necessary for soldier development in the ant *Pheidole hyatti*

**DOI:** 10.1101/2021.03.05.434146

**Authors:** Sophie Koch, Rui Tahara, Angelly Vasquez-Correa, Ehab Abouheif

**Affiliations:** Department of Biology, McGill University, 1205 Avenue Docteur Penfield, Montréal, QC, Canada H3A 1B1

**Keywords:** Inter-organ Communication, rudiments, ecoevo-devo, novel ant castes, *Pheidole*, nano-CT, imaginal disc, wing disc, ant head, ants

## Abstract

The growth of imaginal discs in holometabolous insects is coordinated with larval growth to ensure the symmetrical and proportional development of the adult appendages. In ants, the differential growth of these discs generates distinct castes – the winged male and queen castes and the wingless worker caste. In the hyperdiverse ant genus *Pheidole*, the worker caste is composed of two morphologically distinct subcastes: small minor workers and larger, big-headed soldiers. Although these worker subcastes are completely wingless, soldier larvae develop rudimentary forewing discs that are necessary for generating the disproportionate head-to-body scaling of the big-headed soldier subcaste. However, it remains unclear whether rudimentary forewing discs in soldier larvae are coordinated with other imaginal discs, and whether disc growth and coordination patterns vary between the minor worker and soldier subcastes. Here we show, using quantitative nano-CT three-dimensional analyses, that growth of the soldier rudimentary forewing discs is coordinated with the increase in volume of the leg and eye-antennal (head) discs as well as with larval size. We found that the growth rate of the rudimentary forewing discs differs from the leg discs but is similar to the growth of the head (eye-antennal) discs relative to larval size, suggesting that growth of each type of imaginal disc may be differentially regulated. In addition to their larger size, the soldier eye-antennal discs increase in width as they undergo morphogenesis to generate the characteristic shape of the large soldier head, suggesting that the rudimentary forewing discs may regulate their patterning in addition to their growth. Finally, we observe little growth of the leg and eye-antennal discs during the bipotential stage, while in minor worker development these discs grow at similar rates to one another in coordination with larval size to generate the smaller minor worker subcaste. Our results suggest that rudimentary organs with regulatory functions may participate in new patterns of inter-organ coordination and regulation to produce novel phenotypes and complex worker caste systems. We provide characterization of larval development and imaginal disc growth and morphogenesis with the aim of highlighting this as an emerging system for the study of rudimentary organs during development and evolution.

## Introduction

During development, each organ is specified and undergoes growth and differentiation – a process that is intrinsically regulated, yet also coordinated, with that of other organs and with the growth of the whole organism (Andersen et al., 2013; Droujinine & Perrimon, 2016). This ‘inter-organ coordination’ is phylogenetically widespread and has been especially well documented in the holometabolous insects, which includes flies, butterflies, beetles, and ants (Andersen et al., 2013; Engel, 2015; Droujinine & Perrimon, 2016; Busse et al., 2018; Gontijo & Garelli, 2018; Rosello-Diez et al., 2018; Koch & Abouheif, 2020). Holometabolous insects have radiated into many ecological niches and their success is thought to be in part due to the evolution of discrete ontogenetic life stages, in which embryos pass through a distinct larval stage before they undergo metamorphosis and transition into their adult form (Truman & Riddiford, 1999; Yang, 2001). During this larval stage, the presumptive appendages, such as the legs, wings, head structures (including the eyes and antennae), as well as the genital structures, develop from cell populations called imaginal discs (Held, 2002). Imaginal discs are specified during embryogenesis and, during larval development, they proliferate and are patterned by largely conserved gene regulatory networks (Held, 2002). As the larva transitions to the pupal stage and begins the process of metamorphosis, each disc, and its surrounding peripodial membrane, undergo morphogenesis to develop into the specified appendage (Haynie & Bryant, 1986; Fristrom, 1993). Each disc develops autonomously and behaves like a quasi-independent developmental module, yet at the same time, its growth is regulated to coordinate with other discs and with development of the whole larva (Held, 2002; West-Eberhard, 2003; Andersen et al., 2013). Mechanisms that regulate disc growth and coordination maintain developmental robustness and can also be targeted by selection to generate novel morphologies and scaling relationships (Williams & Carroll, 1993; Emlen & Nijhout, 2000; Emlen et al., 2012; Vallejo et al., 2015; Shingleton & Frankino, 2018).

Ants are holometabolous insects that have evolved colonies with a morphologically distinct winged queen caste, which primarily performs reproductive tasks, and a wingless worker caste, which primarily performs all other tasks such as foraging, brood-care, and defense (Hölldobler & Wilson, 2009). Queen and worker castes are polyphenic, which means that they are determined during development by environmental cues such as nutrition, temperature, and social interactions (Brian, 1963; Penick & Liebig, 2012; Lillico-Ouachour & Abouheif, 2017). In some lineages, the worker caste has evolved inter-individual variation in size and head-to-body size allometry (disproportionate scaling) called ‘worker polymorphism’ (Wilson, 1953; Fjerdingstad & Crozier, 2006; Wills et al., 2018). There are multiple types of worker polymorphism, where each type is defined by the distribution of size and head-to-body size scaling of individuals within the colony (Wilson, 1953). Ants in the hyperdiverse genus *Pheidole* are a classic example of ‘dimorphic allometry’, where the minor worker and soldier subcastes form two discrete allometric lines (Wilson, 1953; Wilson, 2003). The smaller minor workers generally perform brood-care, nest maintenance, and basic foraging, while larger soldiers contribute more to defense and efficiently perform specific foraging tasks, such as seed-milling (Wilson, 1984; Freener, 1987; Powell & Clark, 2004; Pfeiffer et al., 2006; Mertl & Traniello, 2009; Wills et al., 2018). Many studies have examined the morphological, behavioral, and ecological function, as well as the evolvability, of adult morphology in *Pheidole* worker subcastes, yet much remains to be explored as to how their morphology is generated during development (Wilson, 1984; Pie & Traniello, 2007; Economo et al., 2015; Holley et al., 2016).

Differentiation of worker subcaste morphology is thought to occur through differential growth of imaginal discs (Wilson, 1953). Although adult workers are wingless, worker larvae often develop wing rudiments that vary in their size, shape, and expression of the wing gene regulatory network (Dewitz, 1878; Wheeler & Nijhout, 1981a; Abouheif & Wray, 2002; Bowsher et al., 2007; Shbailat & Abouheif, 2013). In the hyperdiverse genus *Pheidole*, rudimentary wing disc growth is not activated in minor worker larvae (Figure 1d), while in soldier larvae rudiments of the forewing discs are activated, grow rapidly, and increase together with larval size (Figure 7e) (Wheeler & Nijhout, 1981a; Abouheif & Wray, 2002; Sameshima et al., 2004b; Shbailat & Abouheif, 2013). In comparison, male and queen larvae develop large fore-and hindwing discs that develop into wings used for mating flights (Figure 1b, c) (Abouheif & Wray, 2002; Keller et al., 2014). While the queen-worker developmental switch occurs during embryogenesis, the soldier-minor worker switch occurs during the late larval stage (Figure 1a) (Passera, 1974; Passera & Suzzoni, 1978; Wheeler & Nijhout, 1981b). Worker larvae before this switch are ‘bipotential’ and can develop into soldiers or minor workers (Wheeler & Nijhout, 1983; Wheeler, 1986). During this critical window, a high protein diet activates juvenile hormone (JH), which activates soldier development, while if JH fails to surpass its threshold, minor worker development proceeds (Figure 1a) (Passera, 1974; Wheeler & Nijhout, 1981b, 1983; Rajakumar et al., 2012; Metzl et al., 2018). Soldier rudimentary forewing discs appear following worker subcaste determination in the last larval instar and grow positively and rapidly relative to larval length (Wheeler & Nijhout, 1981a). However, unlike in queen larvae where leg and wing discs undergo morphogenesis during the larval-to-pupal transition, the growth of the soldier rudimentary forewing discs appears to slow towards the end of the larval stage, becoming uncoupled from the leg discs that begin to undergo morphogenesis (Wheeler & Nijhout, 1981a; Shbailat et al., 2010). These rudiments then degenerate during the larval-to-pupal transition to form small pupal wing buds, which disappear before the adult stage (Wheeler & Nijhout, 1981a; Sameshima et al., 2004a; Sameshima et al., 2004b; Shbailat et al., 2010).

**Figure 1.**
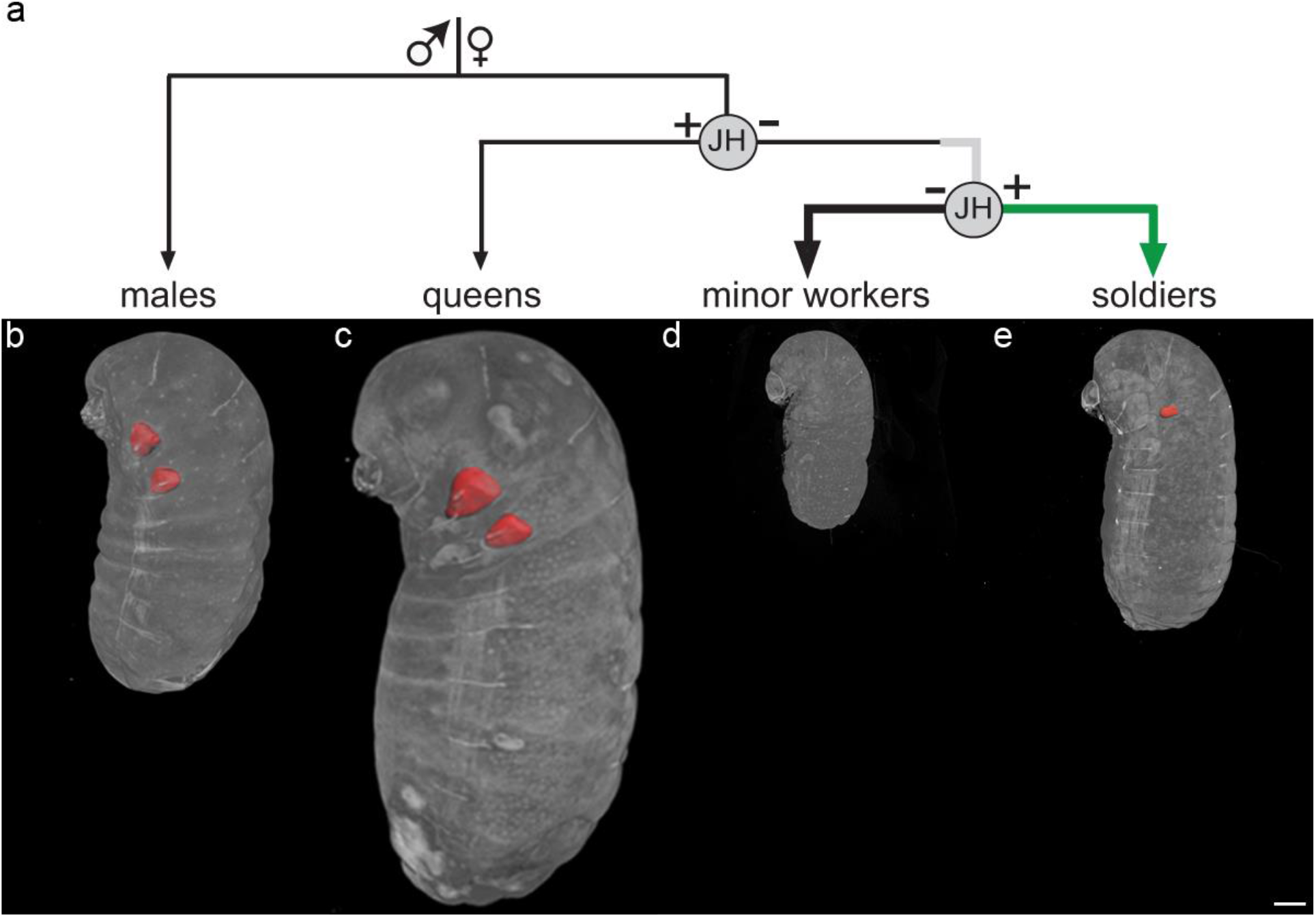
Development and caste determination in *Pheidole*. a) Castes are determined by three developmental switches: 1) the male-female switch is determined by haplo-diploidy during oogenesis, 2) the queen-worker switch during embryogenesis is determined by JH, and 3) the soldier-minor worker switch during larval development is determined by nutrition and JH. Thick grey line represents the bipotential stage, thick black line represents minor worker development, and thick green line represents soldier development. Representative 3D rendered nano-CT images of larvae with fore-and hindwing or rudimentary forewing discs highlighted (red) of b) male, c) queen, and e) soldier. Scale bar indicates 300µm.

Even though the rudimentary forewing disc shows remarkable evidence of regulated growth, gene expression, and apoptosis, their appearance and growth have long been thought to occur due to pleiotropic signaling of JH that activates portions of the queen developmental program (Wheeler & Nijhout, 1981a; Molet et al. 2012, Londe et al.2015, Trible & Kronauer, 2017; Trible & Kronauer 2020). However, Rajakumar et al. (2018) showed that the appearance of rudimentary forewing discs is not a side-effect, but rather, they function to generate size and disproportionate head-to-body scaling of the soldier subcaste (Figure 2g, i). Molecular and physical perturbation of the growth of rudimentary forewing discs generates ‘intermediates’ between wild-type minor workers and soldiers that have altered disproportionate head-to-body scaling. This shows that the growth of the rudimentary forewing discs positively regulates head and body size and is necessary for the development of disproportionate head-to-body scaling in soldiers (Rajakumar et al. 2018). It also suggests that they may participate in a system of inter-organ regulation. However, Rajakumar et al. (2018) results differ from previous observations where perturbations to disc growth result in (1) no detectable adult phenotype due to homeostatic regulation between the damaged disc and other developing discs; or in (2) an increase in the mass of the other adult organs due to differential resource allocation and compensatory growth (Klingenberg & Nijhout, 1998; Nijhout & Emlen, 1998; Parker & Shingleton, 2011; Andersen et al., 2013). Therefore, the regulation between the rudimentary forewing and other imaginal discs in the genus *Pheidole* may be a unique or novel system of inter-organ coordination through which the rudimentary forewing discs generate disproportionate head-to-body allometry (Rajakumar et al. 2018).

**Figure 2.**
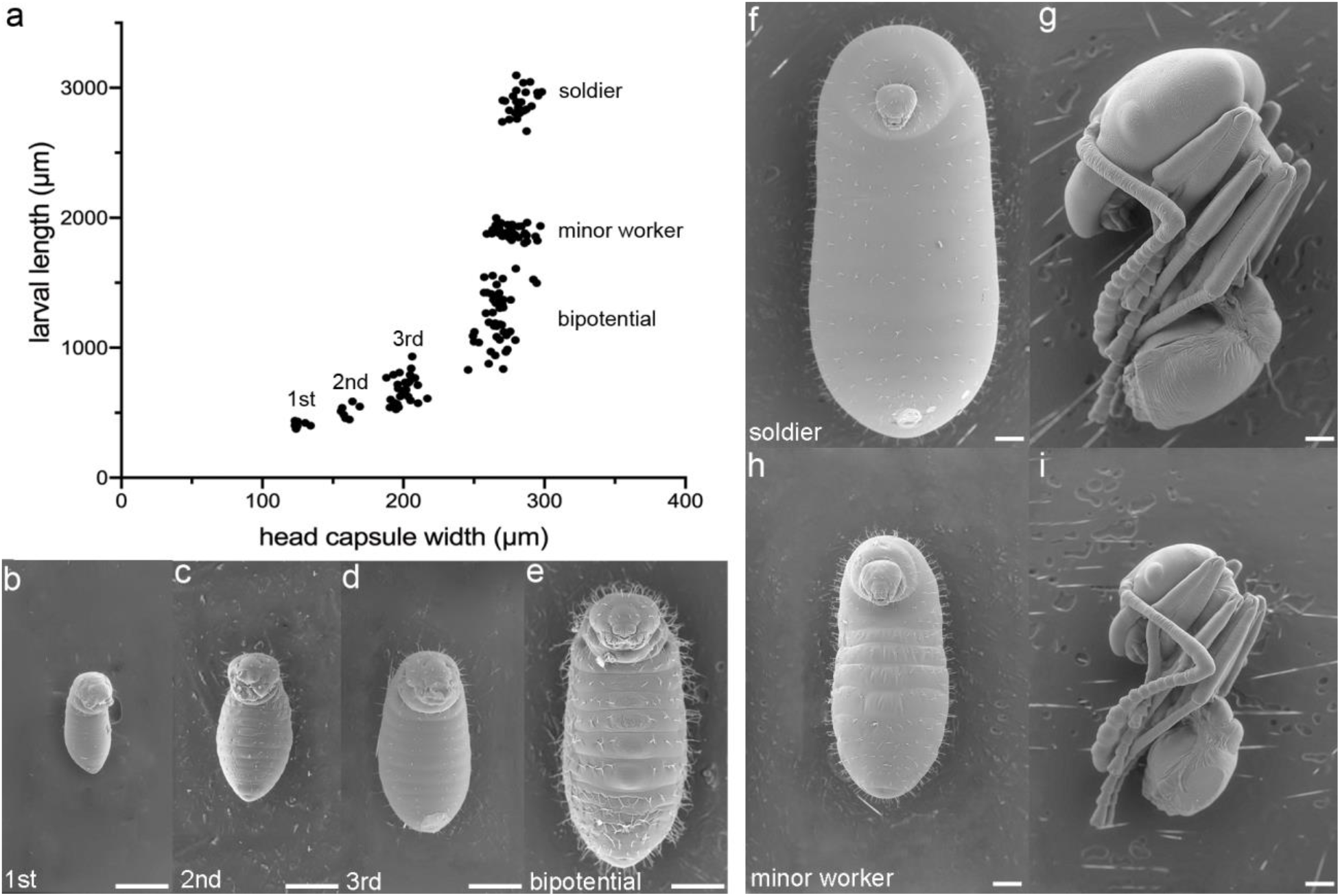
Larval development in *P. hyatti* generates discrete worker subcastes. a) Larval capsule width (µm) versus larval length (µm) identifies four larval instars and differentiation of worker subcastes (n =154). b-e, f, h) Representative individuals of the first (b), second (c), third (d), and fourth (e, f, h) instars in ventral view. The fourth instar includes bipotential larvae (e), minor worker (h), and soldier (f) larvae. Minor worker and soldier larvae represent terminal larvae immediately before metamorphosis. g, i: Representative pupal samples of a soldier (g) and minor worker (i) in left lateral view. Scale bars indicate 200µm.

The main goal of this study is to examine and characterize how the rudimentary forewing discs in soldiers participate in a potentially novel system of inter-organ coordination and how disc growth and coordination vary across worker subcastes. Therefore, in *Pheidole hyatti*, we characterized how the growth of rudimentary forewing discs in soldiers correlates with the growth of the leg and head (eye-antennal) discs within a larva to generate disproportionate head-to-body scaling and how the leg and eye-antennal discs grow and are coordinated with larval size during the bipotential and minor worker stages. The eye-antennal discs give rise to much of the adult head, which differs significantly in size and shape between worker subcastes (Schoeller, 1964; Garcia-Bellido & Merriam, 1969; Milner & Haynie, 1979; Haynie & Bryant, 1986; Wilson, 2003). To investigate these questions, we used X-ray nano-computed tomography (nano-CT) to reconstruct the leg, eye-antennal, and rudimentary forewing discs throughout late larval development. Unlike dissection preparations, nano-CT permits us to (1) characterize multiple organs in whole undissected larva; (2) characterize organs, such as the eye-antennal discs, in three dimensions (3D) because their shape and location render them difficult to quantify in two dimensions (2D); and (3) characterize organs immediately before the larval-to-pupal transition when discs begin to undergo morphogenesis and detach from the larval cuticle (Hall et al., 2017; Watanabe et al., 2017; Schoborg et al., 2018). First, we determined the timing of subcaste determination during larval development and the number of larval instars. We then used nano-CT to examine the growth of the imaginal discs prior to and following the soldier–minor worker developmental switch. The characterization of imaginal disc development by nano-CT will shed light on how a rudimentary organ – the rudimentary forewing discs in *Pheidole hyatti* is coordinated with the growth of the leg and eye-antennal imaginal discs during larval development to generate disproportionate head-to-body scaling and how differential disc growth generates complex worker caste systems.

## Materials and Methods

### Ant collection and care

Queen-right colonies of *Pheidole hyatti* were collected near Globe, Arizona, USA. Colonies were housed at 27°C and 60% humidity on a 12 hour light:dark cycle in the Conviron environmental chambers at the McGill University Phytotron. Nests were constructed from fluon-coated plastic bins with several water-filled glass tubes and a single sugar-water filled glass tube plugged with cotton. Queen-right colonies were fed Bhatkar-Whitcomb yellow diet (Bhatkar & Whitcomb, 1970) and frozen mealworms 3X a week and were supplemented with frozen *Drosophila hydei* 1-2X a week.

### Larval development and scanning electron microscopy (SEM)

To quantify the number of larval instars, larvae were randomly collected from several established colonies and imaged on their dorsum using a Zeiss Discovery V12 stereomicroscope. Larval head capsule width (from the lateral edges of the occipital border) and larval length were measured. These were chosen as morphological landmarks as they have previously been shown to identify: (1) changes in larval head capsule width and larval length associated with larval molts, and (2) changes in larval length associated with worker polymorphism (Wheeler & Nijhout, 1981a; Solis et al., 2010; Alvarado et al., 2015). Larvae designated ‘minor worker’ (Figure 2a, f, g) or ‘soldier’ (Figure 2a, h, i) are terminal larvae immediately before the larval-to-pupal transition are characterized by a dark brown-black gut and changes in fat distribution (Wheeler & Wheeler, 1976). Before this stage, larvae in the first to third instar and the early fourth instar have a light brown gut (Passera, 1974). During the fourth instar, the gut becomes darker brown until it blackens immediately before the prepupal stage. We use larval length as a proxy for the developmental stage and maturation of the larva within this instar (Passera, 1974). All measurements were performed using Zeiss AxioVision Software v4.9.1. Scanning electron microscopy was done on a Hitachi TM3030 Plus TableTop Scanning Electron Microscope. Larval and pupal scans were performed on live specimens.

### Nano-Computed Tomography (nano-CT) scanning and reconstruction

Larval samples were collected from a single established colony, measured for larval length, and samples were fixed for imaginal disc characterization. As nano-CT analyses are costly and time-intensive, we were limited in our assays to 3 brown gut bipotential larvae (Figure 1a, grey line), 2 brown gut minor worker-destined larvae (Figure 1a, black line), 7 brown gut soldier-destined larvae (Figure 1a, green line), and one black gut terminal larvae of each subcaste just before the larval-to-pupal transition (Table 1). Larvae were assigned to stages and subcastes based on larval length, morphology, and gut colour (Figure 2a) (Passera, 1974; Wheeler & Wheeler, 1976). Minor worker and soldier prepupae were fixed in the same way as larvae. All samples were rehydrated to PBS-Tw(0.1%), dehydrated to 70% EtOH, and then stained for 4-21 days in 1% phosphotungstic acid (PTA; Abcam, ab146206-25G) in 70% EtOH depending on the size of the sample (Metscher, 2009a, b). A small hole was punctured through the cuticle of a representative queen and male samples towards the posterior of the larvae to prevent shrinking during staining. Samples were embedded either in 0.5% agarose in a microcentrifuge tube or in 70 % EtOH in a 200µl pipette tip. Images were acquired on a Zeiss Xradia 520 Versa (Carl Zeiss Canada Ltd., ON, Canada) with a resolution ranging from 0.7µm to 3.6µm using 4X or 20X objective lens and single FOV mode. The source and secondary filters were chosen based on the sample. 3201 projections were taken for each specimen over 360 degree-scan. Scans were focused on the anterior half of the larvae to visualize the anterior imaginal discs at high resolution. Detailed information of sample preparation and image acquisition parameters is in Table 1. All three-dimensional reconstructions were done on Imaris x64 v9.2.1 (Bitplane AG, Switzerland). Individual imaginal discs, including their peripodial membrane, and larval brain were segmented and surfaces were filtered manually using 2-6µm resolution to ensure that the disc and peripodial membrane were included, while other larval tissues were excluded in the surface. Volumes were calculated using the Imaris statistics tool. In regions where adjacent discs began to fuse, the surface cutting tool was used to separate them. Prepupal samples were segmented as a single 3D surface then false coloured to capture the entirety of their morphology.

**Table 1.**
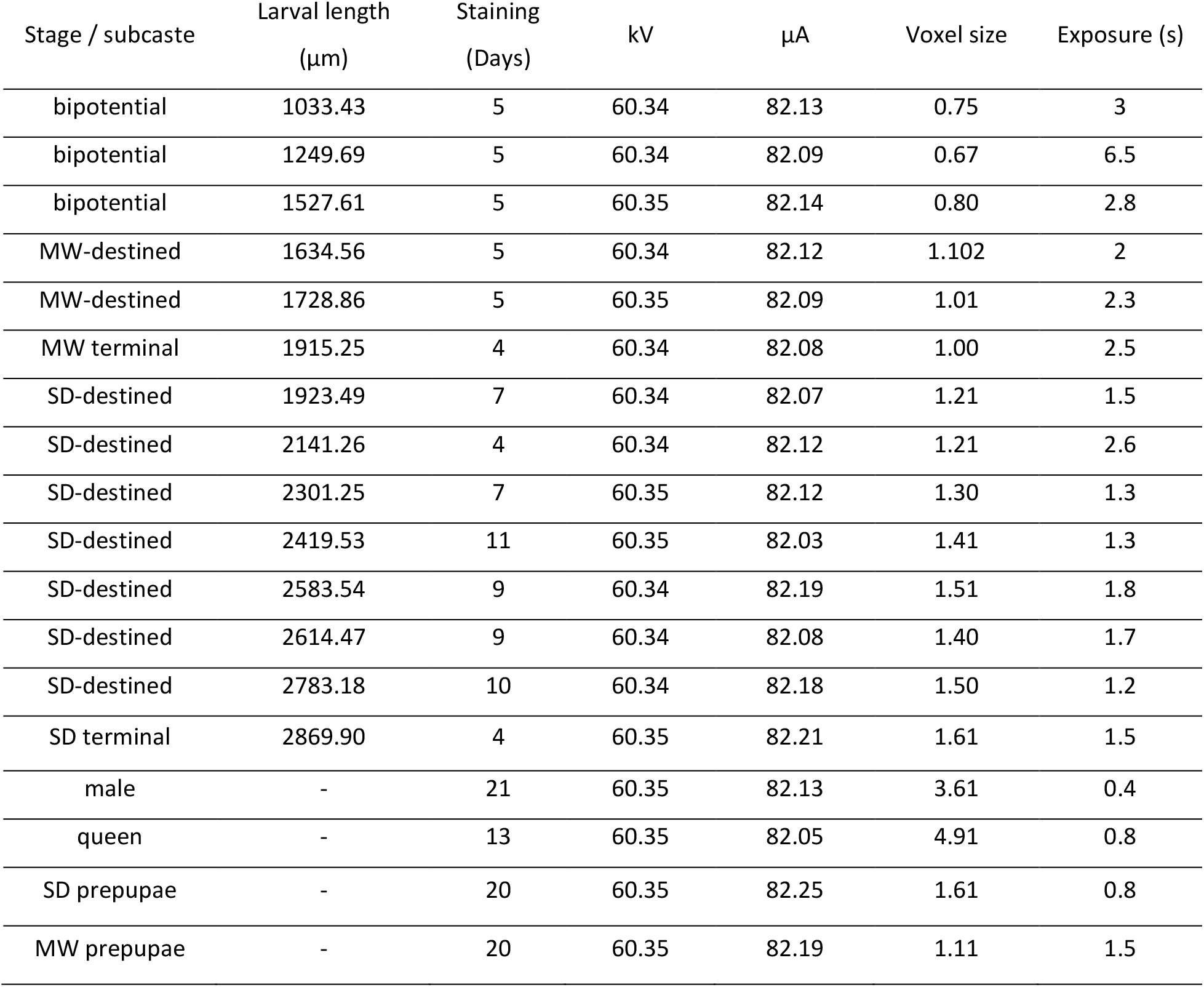
Larval and prepupal sample preparation and image acquisition parameters for nano-CT scans. MW = minor worker, SD = soldier

| Stage / subcaste | Larval length<br>($\mu\text{m}$) | Staining<br>(Days) | kV | $\mu\text{A}$ | Voxel size | Exposure (s) |
| --- | --- | --- | --- | --- | --- | --- |
| bipotential | 1033.43 | 5 | 60.34 | 82.13 | 0.75 | 3 |
| bipotential | 1249.69 | 5 | 60.34 | 82.09 | 0.67 | 6.5 |
| bipotential | 1527.61 | 5 | 60.35 | 82.14 | 0.80 | 2.8 |
| MW-destined | 1634.56 | 5 | 60.34 | 82.12 | 1.102 | 2 |
| MW-destined | 1728.86 | 5 | 60.35 | 82.09 | 1.01 | 2.3 |
| MW terminal | 1915.25 | 4 | 60.34 | 82.08 | 1.00 | 2.5 |
| SD-destined | 1923.49 | 7 | 60.34 | 82.07 | 1.21 | 1.5 |
| SD-destined | 2141.26 | 4 | 60.34 | 82.12 | 1.21 | 2.6 |
| SD-destined | 2301.25 | 7 | 60.35 | 82.12 | 1.30 | 1.3 |
| SD-destined | 2419.53 | 11 | 60.35 | 82.03 | 1.41 | 1.3 |
| SD-destined | 2583.54 | 9 | 60.34 | 82.19 | 1.51 | 1.8 |
| SD-destined | 2614.47 | 9 | 60.34 | 82.08 | 1.40 | 1.7 |
| SD-destined | 2783.18 | 10 | 60.34 | 82.18 | 1.50 | 1.2 |
| SD terminal | 2869.90 | 4 | 60.35 | 82.21 | 1.61 | 1.5 |
| male | - | 21 | 60.35 | 82.13 | 3.61 | 0.4 |
| queen | - | 13 | 60.35 | 82.05 | 4.91 | 0.8 |
| SD prepupae | - | 20 | 60.35 | 82.25 | 1.61 | 0.8 |
| MW prepupae | - | 20 | 60.35 | 82.19 | 1.11 | 1.5 |

### Soldier rudimentary forewing disc area measurements

Soldier larvae were collected randomly from several established lab colonies based on larval size and gut colour characteristics. Larvae were imaged for larval development characterization and measured for larval length, then fixed for immunohistochemistry and counter-stained with DAPI before flat mounting in 50% glycerol. Disc area measurements were performed to exclude the peripodial membrane on a Zeiss AxioImager Z1 microscope using Zeiss AxioVision software v4.9.1. Measurements were eliminated due to disc tearing or poor boundary visibility.

### Statistics

All statistical analyses were performed in Prism v8 and we considered P<0.05 statistically significant. For multiple comparisons, our level of significance was corrected using a Bonferroni correction. All measures of area and volume were log-transformed before analysis (untransformed volumetric estimations can be found in Figure 8). ANCOVA analysis was used to compare the slopes of linear regressions for average imaginal disc area or volume relative to larval length or leg disc volume. If no difference in slope was detected, ANCOVA was used to compare the y-intercepts. We note that our small and unequal sample size between subcastes puts us at risk of Type 2 errors, therefore any non-significant results were interpreted with caution.

## Results

### Larval development and worker subcaste differentiation

To characterize and identify the number of larval instars and size variation within them, we measured larvae for larval head capsule and larval length (Solis et al., 2010). In *P. hyatti*, these measurements identified four larval instars (Figure 2a). The first instar ranges from 376-438µm in larval length, the second instar ranges from 448-586µm, the third instar ranges from 527- 932µm, and the fourth (final) instar, which shows significant variation in larval length including bipotential, minor worker, and soldier larvae, ranges from 830-3096µm (Figure 2a-f, h). (Figure 2a). The younger larvae in this instar are ‘bipotential’ (Figure 1a, gray line, and 2e) and have the capacity to develop either into soldiers (Figure 1f) or minor workers (Figure 1h). Following the JH-mediated developmental switch, the minor worker and soldier developmental programs proceed and differentiate. As larvae approach the end of larval development, the light brown gut continues to darken until the terminal stage, at which time, it becomes dark black, and larvae show changes in fat distribution (Passera, 1974; Wheeler & Wheeler, 1976). Minor worker larvae become terminal between 1804-1998µm (Figure 2h), whereas soldier larvae grow larger and become terminal between 2666-3096µm (Figure 2f). Before the terminal stage, larvae are ‘minor worker-destined’ (Figure 1a, black line) or ‘soldier-destined’ (Figure 1a, green line) and have a light brown gut. Differentiation within the fourth instar produces two discrete clusters of terminal stage larvae (Figure 2a). These terminal clusters together generate the discrete dimorphic head-to-body allometry that is characteristic of typical *Pheidole* worker caste systems (Figure 2g, i).

### Quantitative nano-CT analyses of imaginal disc growth and coordination

We describe the growth and morphogenesis of these imaginal discs and compare the slope and y-intercepts of their linear regressions within subcastes to investigate their growth relative to larval length and relative to leg disc volume.

### Coordination of imaginal disc development in bipotential larvae

In the bipotential stage (830-1600µm in larval length), larvae have the capacity to develop into soldiers or minor workers and can be experimentally induced to a soldier fate by exogenous application of a JH-analog (Wheeler & Nijhout, 1983; Rajakumar et al., 2012). During this stage, the leg discs are spherical (Figure 3) and their volume does not increase significantly relative to larval length (F=1.660, df =1, P=0.4202) (Figure 7a, d). The three pairs of leg discs are positioned ventrally, inferior to the larval head capsule and lateral to the ventral nerve cord. The eye-antennal discs also begin as spherical discs and are positioned anterosuperior to the larval brain within the larval head capsule (Figure 3). Throughout bipotential development, the eye-antennal discs become slightly more oblong and extend anteriorly (Figure 3). During this stage, the volume of the eye-antennal discs does not correlate with larval length (F=4.436, df=1, P=0.2822) but does correlate with leg disc volume (Figure 7b, d). Relative to larval length, the eye-antennal discs do not differ significantly in slope or y-intercept compared to the leg discs (slope: F=0.6733, df=2, P=0.4982; y-intercept: F=7.104, df=3, P=0.0760) (Figure 7d) suggesting that the slight growth of the leg and eye-antennal discs occurs at a similar rate during the bipotential stage but does not correlate with larval length. We were unable to detect rudimentary forewing or hindwing discs in bipotential samples (Figure 3).

**Figure 3.**
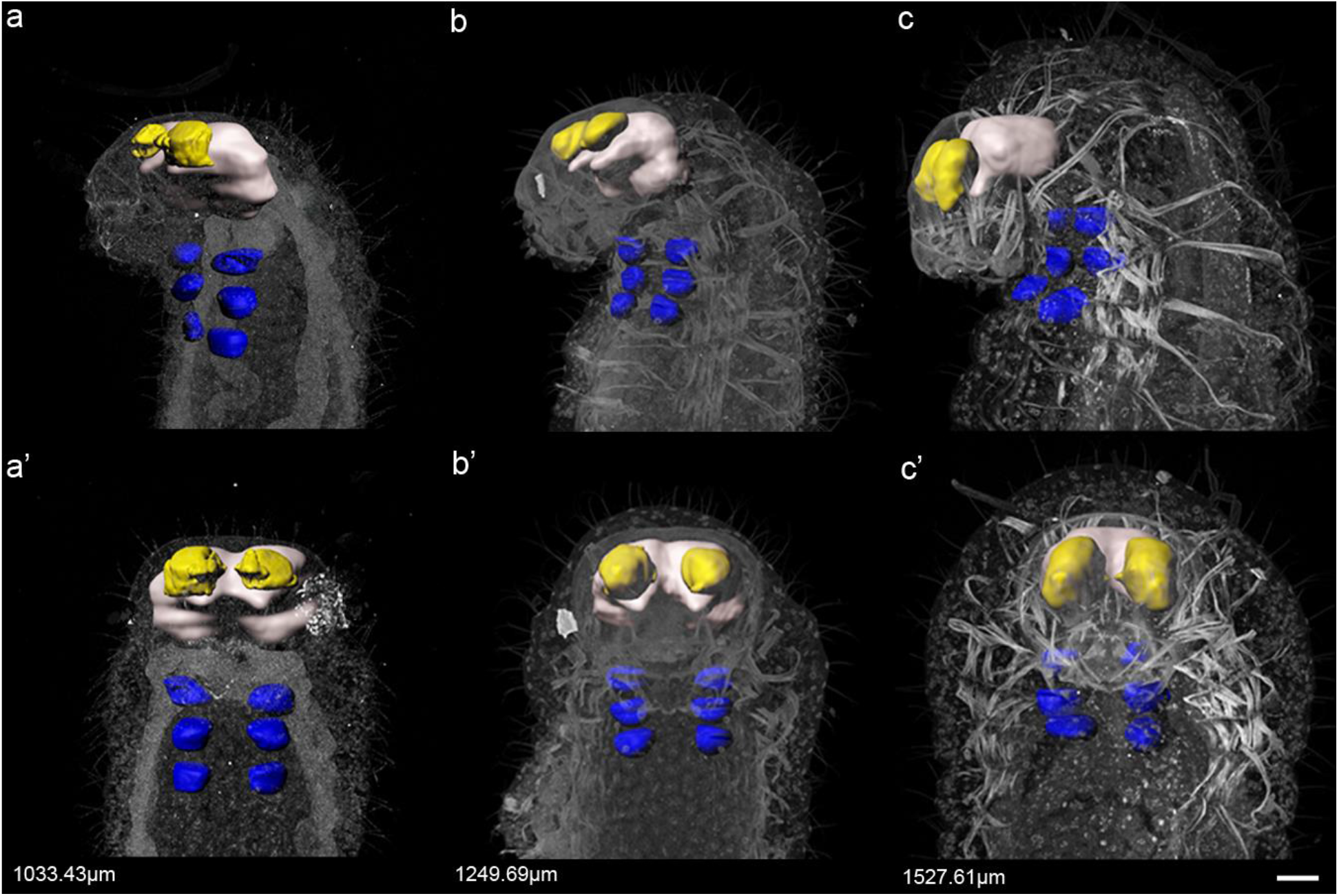
3D nano-CT analysis of imaginal disc development in bipotential larvae. Left lateral (a-c) and ventral (a’-c’) views of 3D nano-CT reconstructions of bipotential larvae: 1033.43µm (a, a’), 1249.69 (b, b’), and 1527.61µm (c, c’) in length. Blue represents leg imaginal discs, yellow represents eye-antennal discs, and white represents the larval brain. Larval length in µm is indicated in the bottom left corner. Rudimentary forewing discs are undetectable. Images to scale, scale bar = 70µm.

### Coordination of imaginal disc development in minor worker larvae

During minor worker development (1600-1998µm in larval length), the leg discs increase positively in volume relative to larval length (Figure 7a, d). In terminal larvae, the legs begin to undergo morphogenesis, increase significantly in volume, and their peripodial membranes begin to evert and merge with those of the adjacent discs (Figure 4c, c’). Similar to the leg discs, the eye-antennal discs increase positively in volume relative to larval length (Figure 7b, d) and increase positively relative to leg disc volume (Figure 7e). The eye-antennal discs become more oblong, extending anteriorly into the larval head capsule as minor worker development progresses (Figure 4a, b, a’, b’). The increase in volume of the eye-antennal discs continues until, at the end of minor worker larval development, they undergo morphogenesis: the presumptive antenna evert, and the peripodial membranes fuse along the midline, expanding laterally to envelop the larval brain (Figure 4c, c’). In minor worker larvae, eye-antennal discs have a significantly larger y-intercept compared to leg disc volume but do not differ in slope (slope: F=16.15, df=2, P=0.0567; y-intercept: F=22.24, df=3, P=0.0181) (Figure 7d). We observed very small forewing rudiments in one minor worker-destined sample (Figures 4b, b’ and 7c) that were no longer detectable by the terminal stage (Figure 4c, c’).

**Figure 4.**
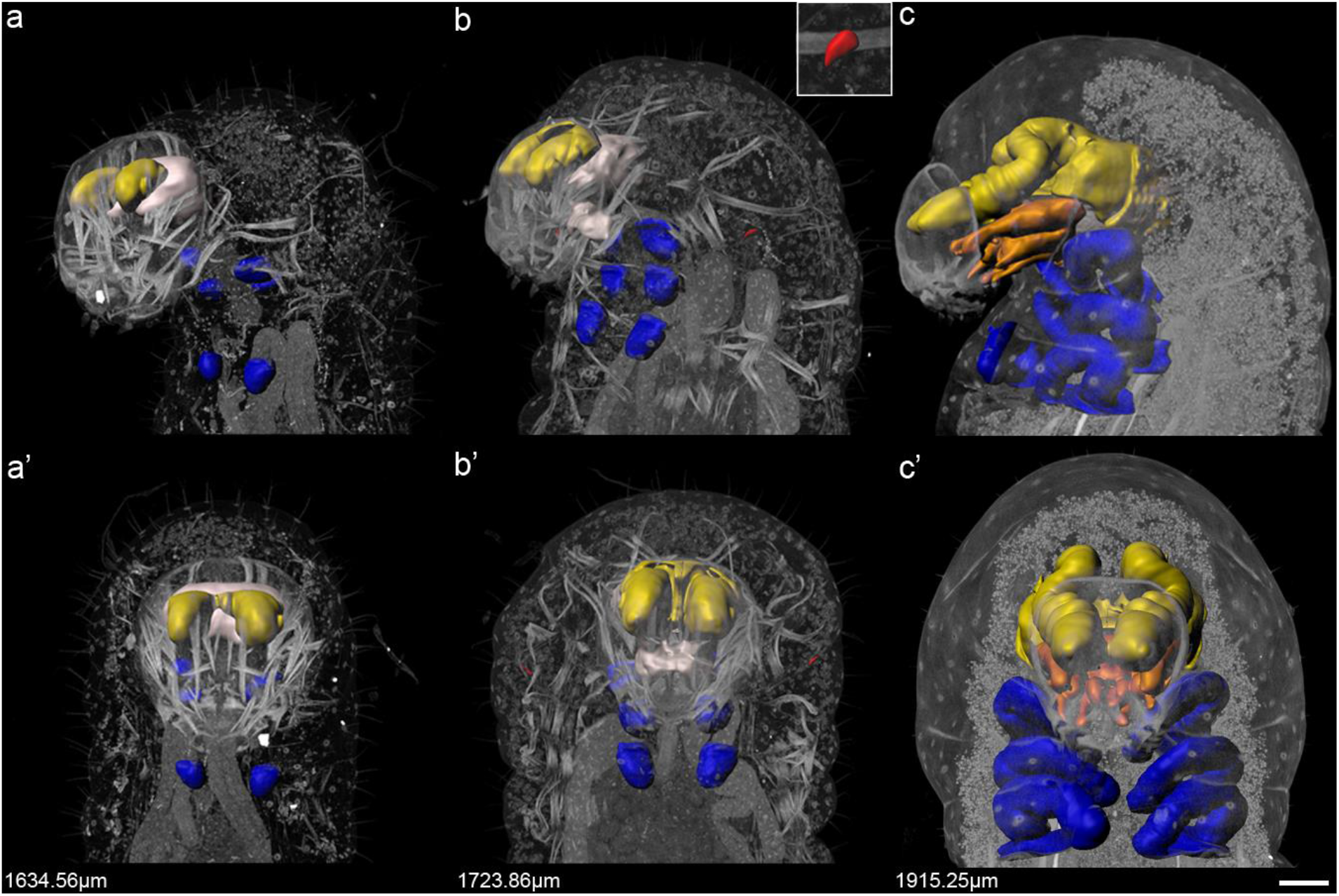
3D nano-CT analysis of imaginal disc development in minor worker larvae. Left lateral (a-c) and ventral (a’-c’) views of 3D nano-CT reconstructions of minor worker-destined (a, a’, b, b’) and terminal minor worker (MW; c, c’) larvae: 1634.56µm (a, a’), 1723.86µm (b, b’), and 1915.25µm (c, c’) in length. Inset in (b) is magnified to show rudimentary forewing discs in sample (b, b’). Larval length in µm is indicated in the bottom left corner. Blue represents leg imaginal discs, yellow represents eye-antennal discs, orange in (c, c’) represents labial and/or clypeo-labral discs, and white represents the larval brain. Note that in (b, b’) the distinct mass labeled white under the main larval brain (also white) may correspond to the labial and/or clypeo-labral disc before its differentiation into clear mouthparts. However, the resolution of our scan was not sufficient to confirm whether this distinct mass labeled white under the main larval brain represents undifferentiated mouthparts and therefore should be labeled orange. Rudimentary forewing discs were undetectable in (a, a’) and (c, c’). Images comparisons to scale, scale bar = 100µm.

We were able to visualize discs near the larval mouthparts in the terminal minor worker sample that appear to be derived from the labial and/or clypeo-labral discs that contribute to the mandibles and other mouthparts (Figure 4c, c’) (Kumar et al., 1979; Held, 2002). At the stage just prior to this (b, b’), the distinct mass (labeled white) under the main larval brain (also labeled white) may correspond to the labial and/or clypeo-labral disc before its differentiation into clear mouthparts. However, the resolution of our scan was not sufficient to confirm whether this distinct mass represents undifferentiated mouthparts. At the terminal stage, these discs then merge with the eye-antennal discs and appear to vary in shape between subcastes (Figure 8a-b’). By the prepupal stage, the mandibles and other mouthparts are more clearly visible (Figure 8c-d’). Based on the prepupal scans that we performed, we estimated the boundary between the presumptive mandibles and other mouthparts from the eye-antennal discs in our terminal samples and then removed the former from our estimation of the eye-antennal discs as they are likely derived from other discs (Figure 8).

### Coordination in imaginal disc development in soldier larvae

In addition to a larger terminal size compared to minor workers, soldier-destined larvae are characterized by the growth of the rudimentary forewing discs (Figure 6). The soldier rudimentary forewing discs differ considerably from the queen and male forewing discs in size and shape, and hindwing rudiments are undetectable (Figure 1b, c, e). To estimate how the rudimentary forewing discs grow relative to the leg discs in soldier larvae, we first measured the area of the discs relative to larval length using standard dissection techniques. The three pairs of leg discs increase in area positively relative to larval length and there is no significant difference in slope or y-intercept between the average disc area across the three thoracic segments (slope: F=0.2961, df=2, 113, P= 0.7443; y-intercept: F=0.3184, df=2, 113, P =0.7280) (Figure 5a). Using nano-CT 3D analyses, we observed that in soldier-destined larvae with approximately the same larval length as the terminal minor worker larvae the leg discs remain spherical (compare Figure 4c and 6a). Throughout the soldier-destined period, they continue to increase positively in volume relative to larval length (Figure 7a, d), acquiring visible segments along their proximo-distal axes (Figure 6a-g, a’-g’). Similar to minor workers, as the soldier larvae approach the larval-pupal transition, the leg discs undergo morphogenesis, increase significantly in volume, and detach from the larval cuticle (Figure 6h, h’). By the end of larval development, the average leg disc volume in the terminal soldier sample is approximately 2.5 times larger than the average leg disc volume of the minor worker sample (Figure 7a).

**Figure 5.**
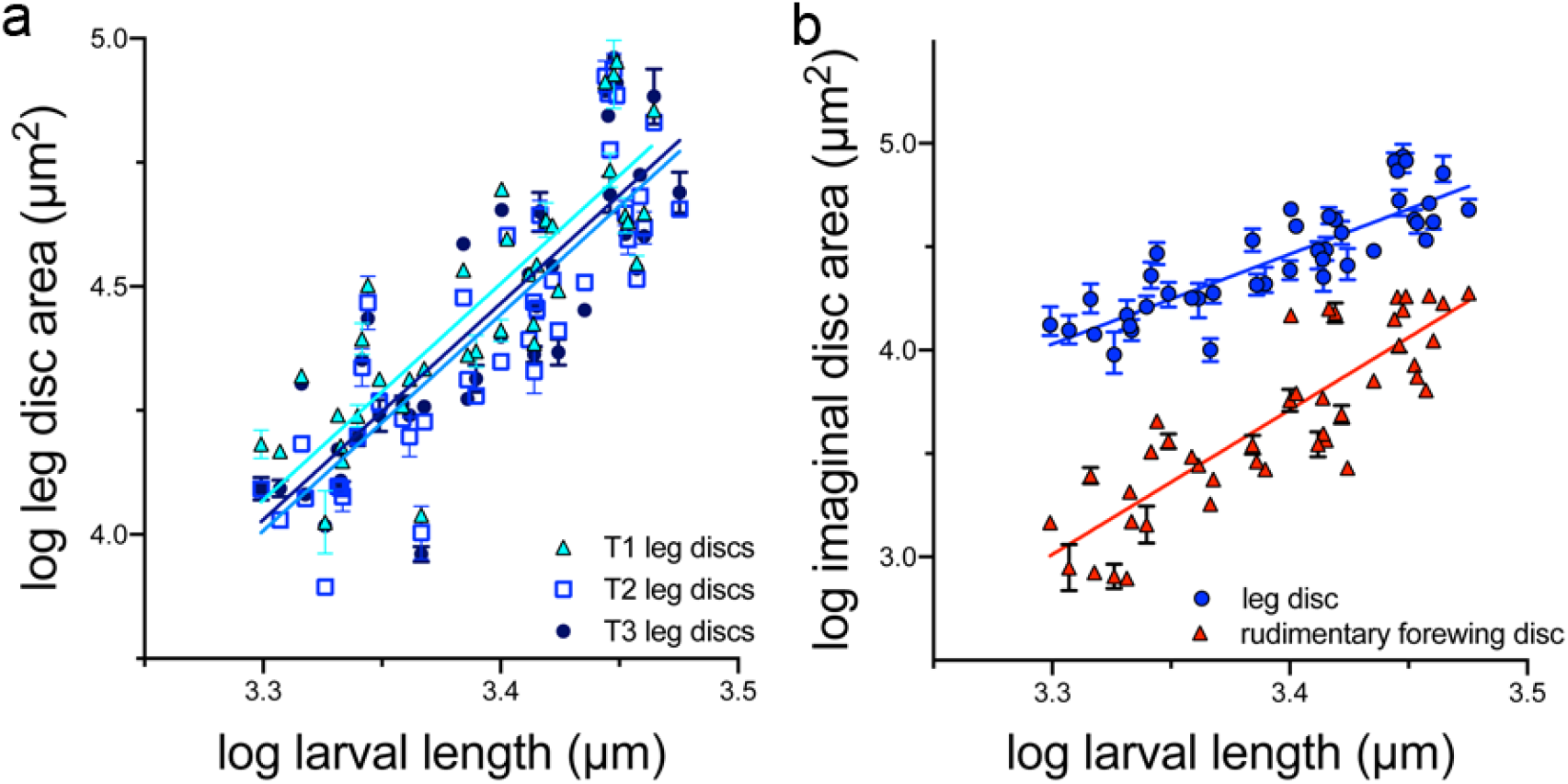
Rudimentary forewing discs grow rapidly during soldier development. Imaginal disc growth during soldier development: a) Linear regressions of log larval length (µm) versus average log leg disc area (µm2) separated based on thoracic segment identity (T1-T3) (n=43 larvae): T1 leg discs = turquoise circles (n=37), T2 leg discs = blue squares (n=41), T3 leg discs = dark blue squares (n=41) (slope: F=0.2961, df=2, 113, P= 0.7443; y-intercept: F=0.3184, df=2, 113, P =0.7280). Best fit by a single curve y = 4.343x – 10.29. b) Linear regressions of log larval length (µm) versus log area (µm2) of soldier leg (blue) and rudimentary forewing (red) discs (n = 43 larvae) (F=11.51, df=82, P=0.011). Symbols indicate mean and error bars indicate range.

**Figure 6.**
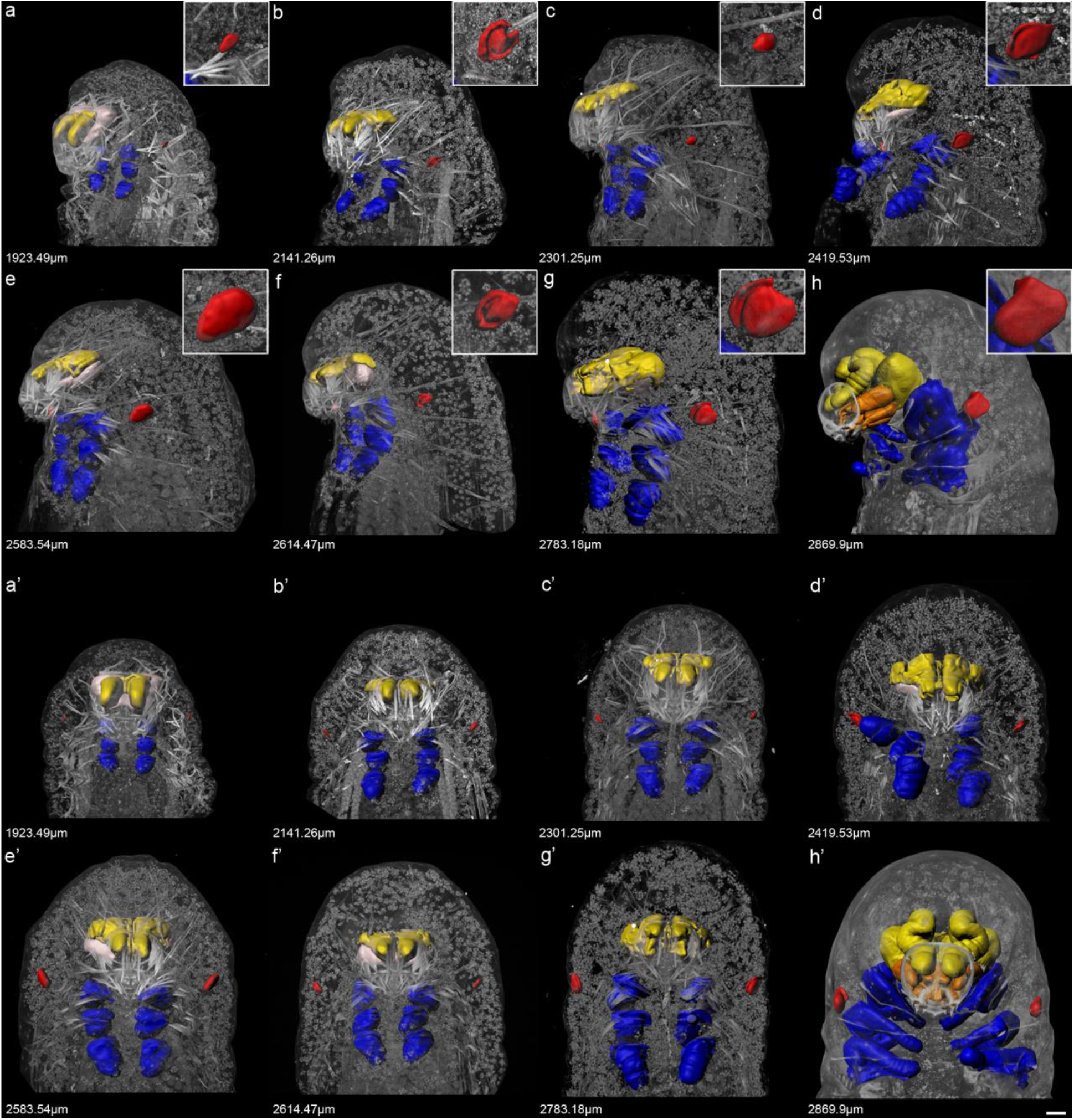
3D nano-CT analysis of imaginal disc development in soldier larvae. Left lateral (a-h) and ventral (a’-h’) views of 3D nano-CT reconstructions of soldier-destined (a-g, a’-g’) and terminal soldier (SD; h, h’) larvae: 1923.49µm (a, a’), 2141.26µm (b, b’), 2301.25µm (c, c’), 2419.53µm (d, d’), 2583.54µm (e, e’), 2614.47µm (f, f’), 2783.18µm (g, g’), and 2869.9µm (h, h’) in length. Insets in a-h show magnified rudimentary forewing discs in the corresponding sample. Larval length in µm is indicated in the bottom left corner. Blue represents leg imaginal discs, yellow represents eye-antennal discs, orange in (h, h’) represents labial and/or clypeo-labral discs, white represents the larval brain, and red represents rudimentary forewing discs. Image comparisons to scale, scale bar = 100µm.

During soldier development, the eye-antennal discs continue to grow in volume and length, extending distally into the larval head capsule (Figure 6a-g, a’-g’). The peripodial membranes that surround the eye-antennal discs begin to fuse along their medial edges until, at the late soldier-destined stage, the membranes further expand laterally around the larval brain (Figure 6g, g’). Throughout soldier development, eye-antennal disc volume correlates positively with larval length and leg disc volume (Figure 7b, d, e). Relative to larval length, the soldier eye-antennal discs are larger and differ significantly in y-intercept compared to the leg discs (F=8.460, df=13, P=0.0122) but do not differ in slope (F=0.03928, df=12, P=0.8462) (Figure 7d). In the terminal soldier larvae, the eye-antennal discs also undergo morphogenesis and are approximately 1.43 times larger in volume than those in minor workers (Figures 6h, h’ and 7b). At the terminal stage, the peripodial membrane expands to surround the larval brain and forms the presumptive adult head. In addition to being larger in volume, the width of the peripodial membrane is larger in soldiers than minor workers and appears to further enlarge during the prepupal stage (Figure 8b, b’, d, d’). Just as in our terminal minor worker sample, we removed the presumptive mandibles and mouthparts from the estimation of the eye-antennal discs in the terminal soldier sample.

**Figure 7.**
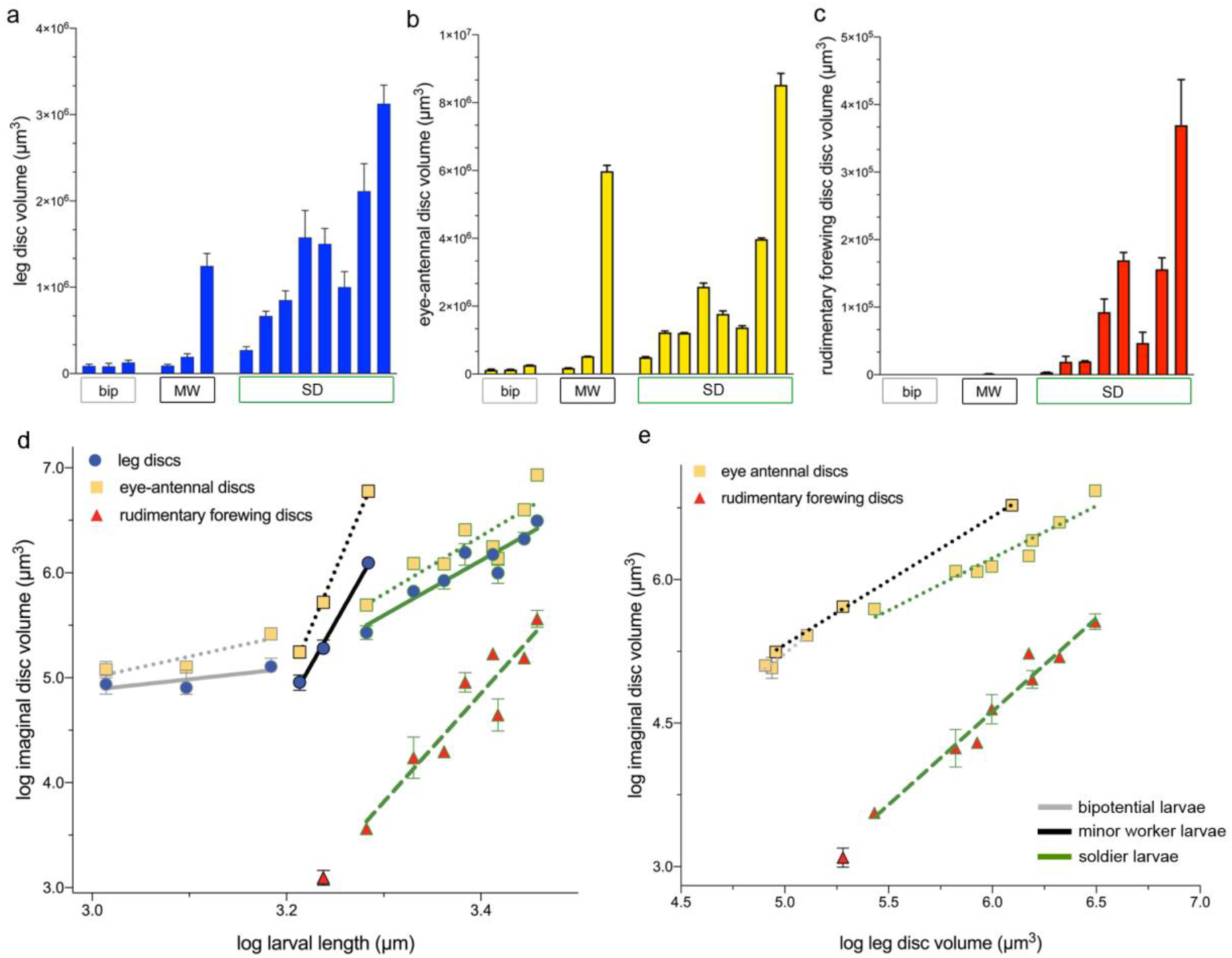
Leg, eye-antennal, and rudimentary forewing disc growth across subcastes. a-c) Bar graphs showing imaginal disc volume (µm3) versus larval length (µm) grouped by developmental stage and subcaste. Bars show mean and error bars indicate range for: (a) leg disc volume (µm3), (b) eye-antennal disc volume (µm3), and (c) rudimentary forewing disc volume (µm3) grouped by bipotential (bip, grey), minor worker (MW, black), and soldier (SD, green) arranged by ascending larval length. d) Linear regressions of log larval length (µm) versus log imaginal disc volume (µm3) grouped by subcaste. e) Linear regression of log leg disc volume (µm3) versus imaginal disc volume (µm3) grouped by subcaste. Each point represents the mean and range within a larva. Blue circles represent leg discs (solid lines), yellow squares represent eye-antennal discs (dotted lines), and red triangles represent rudimentary forewing discs (dashed lines). Grey lines and outlines represent bipotential larvae, black lines and outlines represent minor worker larvae, and green lines and outlines represent soldier larvae. The level of significance was corrected for multiple comparisons using the Bonferroni correction.

**Figure 8.**
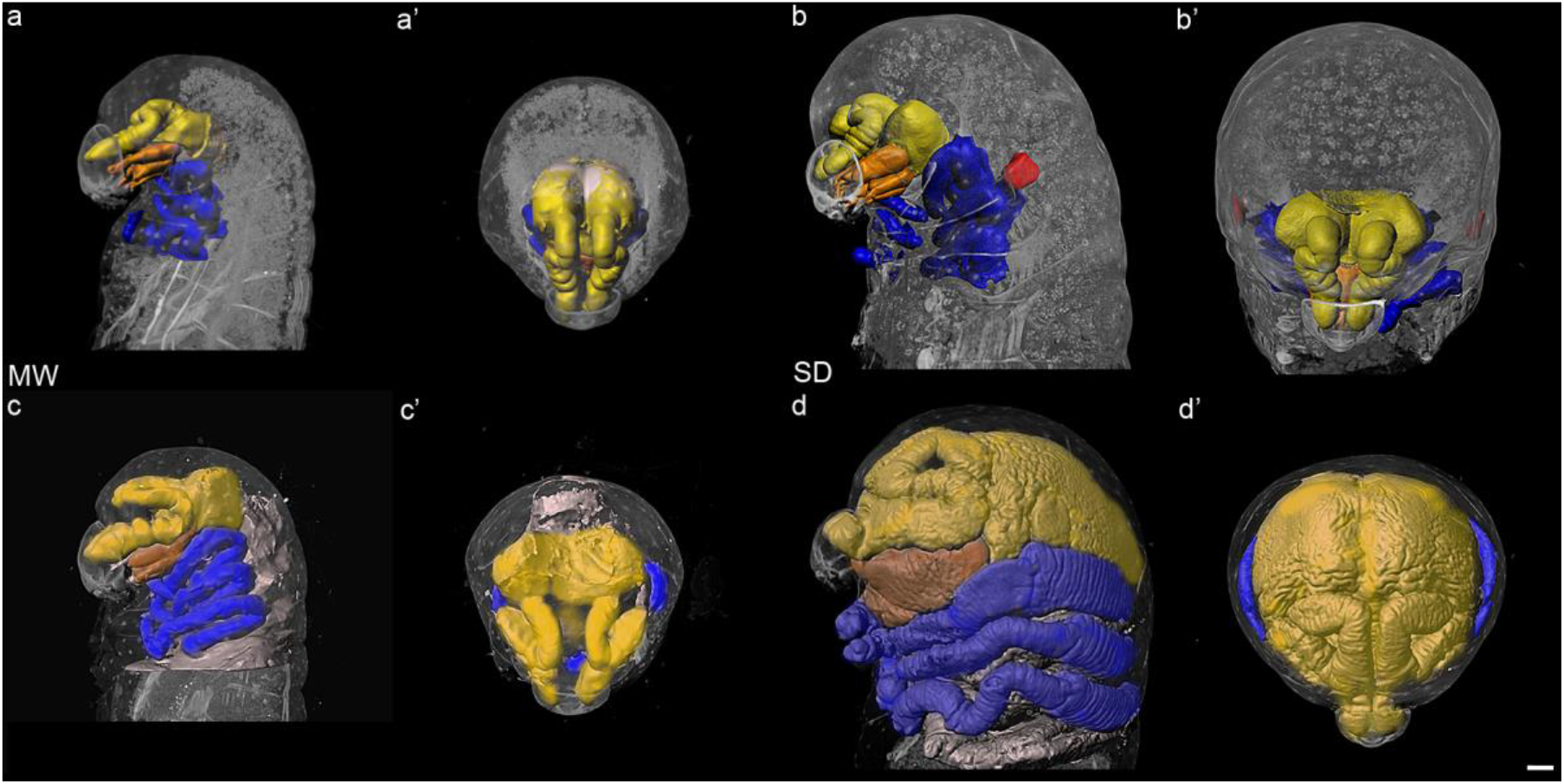
Soldier eye-antennal discs grow larger and generate a wider head during morphogenesis than in minor workers. Left lateral (a-d) and superior (a’-d’) views of 3D nano-CT reconstructions of terminal minor worker (MW) (a, a’), terminal soldier (SD) larvae (b, b’), minor worker prepupae (c, c’), and soldier prepupae (d, d’). Blue represents leg imaginal discs, yellow represents eye-antennal discs, orange represents labial and/or clypeolabral discs, white represents the larval brain in (a-b’), and red represents rudimentary forewing discs. Images to scale, scale bar = 100µm.

The rudimentary forewing discs begin as a small cluster of cells in the early soldier-destined stage and increase in volume rapidly throughout the soldier-destined period, obtaining a more triangular shape in lateral view in late larval development (Figures 6 and 7c). However, there is some plasticity in the overall ontogenetic trajectory between developing soldier larvae (Figure 6e-g). The soldier rudimentary forewing discs generally increase in both area and volume relative to larval length and relative to leg disc volume (Figures 5b and 7). Relative to larval length, the area of soldier rudimentary forewing discs increases at a significantly different rate compared to the average leg disc area (F=11.51, df=82, P=0.011) (Figure 5b) and the volume of the soldier rudimentary forewing discs also increases at a significantly different rate compared to the volume of the soldier leg discs (F=8.806, df=12, P=0.0118). The increase of soldier rudimentary forewing disc volume is not significantly different from the soldier eye-antennal disc volume (slope: F=5.879, df=12, P=0.0320, y-intercept: F=7.053, df=12, P=0.0210) (Figure 7d). However, when plotted relative to leg disc volume, soldier rudimentary forewing disc volume increases at a significantly different rate compared to the soldier eye-antennal discs (F=17.31, df=12, P=0.0013) (Figure 7e). In terminal larvae, the rudimentary forewing discs detach from the larval cuticle and remain attached to the larval fat body through their peripodial membrane (Figure 6h, h’), however, unlike the leg and eye-antennal discs, they do not undergo morphogenesis.

## Discussion

The growth of the soldier rudimentary forewing discs positively regulates the growth of head and body size to generate disproportionate head-to-body scaling and the soldier phenotype in *P. hyatti* (Rajakumar et al. 2018). These results suggest that these rudiments may regulate the differential growth of the other discs through a potentially novel system of inter-organ coordination. Here we describe the growth and development of three imaginal discs and show that the activation of the rudimentary forewing discs is coordinated with an increase in the volume of the leg and eye-antennal discs and with an increase of larval size. Furthermore, as the soldier eye-antennal discs undergo morphogenesis, they increase in width to specify the larger soldier head and generate its characteristic shape. In contrast, little imaginal disc growth occurs during the bipotential stage, whereas the leg and eye-antennal discs increase proportionally in minor worker development. Collectively, this shows stage and subcaste-specific differences in disc growth and morphogenesis. This study provides a foundation for understanding how these rudiments grow and participate in inter-organ coordination to generate disproportionate head-to-body scaling. More generally, we highlight the importance of characterizing development across stages and subcastes to investigate the evolution of complex worker caste morphology in ants.

The developmental differentiation that generates worker polymorphism occurs during the last larval stage (Wheeler & Nijhout, 1981a, 1983; Sameshima et al., 2004b; Alvarado et al., 2015). In *P. hyatti* and other species of *Pheidole*, this differentiation produces discrete clusters of terminal larvae that generate the dimorphic minor workers and soldiers that both develop from the early bipotential stage. This contrasts with the carpenter ant, *Camponotus floridanus*, which has a continuous worker caste system of minors, media, and major workers, and whose fourth instar is represented by early bipotential larvae and a continuous spread of terminal larval sizes (Alvarado et al., 2015). This shows that modifications to larval growth and terminal size can generate different types of worker polymorphism in ants (Wilson, 1954). The developmental mechanisms that underlie the differences and distributions of larval size within a worker caste system involve JH, nutrition-based signals, such as the insulin/insulin-like pathways, social pheromone-based regulation, and/or epigenetic modifications, yet much remains to be explored as to how these signaling pathways interact during larval development and vary between complex caste systems (Wheeler & Nijhout, 1984; Wheeler, 1986; Rajakumar et al., 2012; Alvarado et al., 2015; Corona et al., 2016; Lillico-Ouachour & Abouheif, 2017). While larval length correlates positively with adult body size, it does not entirely explain changes in disproportionate head-to-body scaling (Wilson, 1953; Alvarado et al., 2015). Therefore, we focused our characterization on the differential growth and morphology of the larval imaginal discs, the precursors of the adult structures in holometabolous insects

Our results show that, following soldier subcaste determination, the growth of the rudimentary forewing discs is activated, while the leg and eye-antennal discs continue to grow and reach a final larger size, which is correlated with an increase in larval size. This shows that the growth of the soldier rudimentary forewing discs is coordinated with the other imaginal discs and with larval growth. Within a larva, smaller rudimentary forewing discs co-occur with smaller leg and eye-antennal discs, suggesting that coordination of growth between discs may be more constrained compared to their coordination with overall larval size. Each disc’s growth trajectory may be determined by organ-specific sensitivity to systemic endocrine signals, such as JH or insulin, or due to endogenous properties of the disc (Emlen et al., 2012). The rudimentary forewing discs grow more rapidly compared to the leg discs, which is consistent with ours and previous results using area to estimate disc growth (Wheeler & Nijhout, 1981a). However, we observe that the growth of the rudimentary forewing discs does not differ significantly from the growth of the eye-antennal discs relative to larval length. This pattern may be a result of the intimate relationship between the rudimentary forewing discs and head growth. This is consistent with Rajakumar et al.’s (2018) finding that perturbations to rudimentary forewing disc growth have a more significant effect on pupal head size than body size. However, we cannot rule out that this non-significant relationship between the growth of the rudimentary forewing discs and eye-antennal discs is a result of our low sample size, and thereby, low statistical power. Future studies can investigate this further as well as examine how perturbations to the growth of the solider rudimentary forewing discs affect the growth of the leg and eye-antennal discs.

While the relationship between the leg and rudimentary forewing discs has been previously examined in *P. bicarinata*, the growth of the eye-antennal discs in ant worker subcastes has not been well explored despite the notable phenotypic differences in adult head morphology. We observed that the eye-antennal discs and leg discs increase in volume at similar rates relative to one another during soldier development. However, as the eye-antennal discs undergo morphogenesis, they appear to increase in width as they envelop the larval brain and generate a wide and morphological distinct head cuticle by the prepupal stage. Previous work suggests that modifications to the patterning and the size of the eye-antennal discs may be the source of generating variation in the relative scaling of organs on the head (Dominguez & Casares, 2005). This suggests that the activation of the growth of the soldier rudimentary forewing discs and the soldier developmental program may regulate the increase in eye-antennal disc volume as well as regulate their patterning to specify a larger region of head cuticle and modifications to the antennae and eyes. In contrast, we observed no comparable differences in the leg discs between subcastes other than their size. Therefore, as the soldier rudimentary forewing discs are necessary to produce the solider phenotype, our results suggest that the activation of the soldier developmental program and the rapid growth of the rudimentary forewing discs causes the leg and eye-antennal discs to grow larger throughout soldier development. Furthermore, they may also regulate the patterning of the eye-antennal discs to generate the soldier head shape as the discs undergo morphogenesis.

We observed that during the bipotential stage before worker subcaste differentiation, little disc growth occurs and, while the eye-antennal discs are larger than the leg discs, neither is correlated with larval size. Within our sampled larvae in the minor worker developmental stage, the leg and eye-antennal discs also grow at similar rates but both grow positively relative to larval size. We were unable to detect wing rudiments in our bipotential samples and the majority of our minor worker samples. However, we observed very small wing buds in one sample that we classified as minor worker-destined based on larval size. In addition to the presence of rudimentary forewing discs in soldiers of *P. megacephala*, Sameshima et al. (2004) observed very small populations of forewing cells in minor worker larvae. Unlike in soldiers, these forewing cells in *P. megacephala* are never activated, do not undergo significant growth, and appear to not be coordinated with other discs or with larval growth. These authors also observed forewing cells in a high percentage of larvae smaller than the smallest terminal minor workers, which shows that these cells exist but that their growth is not activated in bipotential or minor worker-destined larvae (Sameshima et al., 2004b). This suggests that either that very small clusters of wing cells develop in the bipotential stage and we were unable to detect them or that Sameshima et al. (2004) also observed these discs in minor worker-destined larvae. Alternatively, this sample may represent a very young soldier-destined larva just after soldier determination. During early destined stages, soldier and minor worker larvae are indistinguishable based on larval characteristics but become distinguishable when the minor worker larval gut begins to darken, while the soldier larval gut remains light and soldier larvae continue to develop. We were unable to resolve these possibilities due to our limited sample size and we acknowledge that our low sample size of bipotential and minor worker-destined larvae make it difficult to conclusively establish the patterns of disc growth and compare statistically between subcastes. However, these results preliminarily show that following worker caste determination, differential growth of imaginal discs and larval development generates discrete worker subcaste morphologies. Future studies can further examine these hypotheses by examining more samples across these stages of development.

Inter-organ coordination appears to be widespread in insects (Koch &Abouheif,2020). Two well-described models have been proposed in the literature: homeostasis and resource competition. According to the inter-organ homeostasis model, the coordination of disc growth functions to ensure developmental symmetry. When a disc is damaged, this regulation coordinates the growth of the other discs with the damaged disc and larval development to allow regeneration, producing no adult phenotype or shows no effects on larval development if the disc is eliminated (Madhavan & Schneiderman, 1969; Parker & Shingleton, 2011; Hariharan, 2012; Gontijo & Garelli, 2018). In contrast, according to the inter-organ resource competition model, damage to one or multiple discs results in an asymmetric increase of the mass of the other appendages suggesting that each disc competes locally for resources (Klingenberg & Nijhout, 1998; Nijhout & Emlen, 1998). In *P. hyatti*, however, perturbations to the growth of the soldier rudimentary forewing discs result in a disproportionate reduction of the growth of the head and body size (Rajakumar et al. 2018). As this differs from previous observations of coordination, this suggests that the soldier rudimentary forewing discs have acquired a novel function. Our characterization of the growth relationships of the leg, eye-antennal, and rudimentary forewing discs supports the hypothesis that they participate in a potentially novel system of inter-organ coordination. While it remains to be explored how these rudiments signal and regulate the growth of the head and body size, previous work on queen and worker castes has shown that imaginal discs growth can be modified to generate caste variation. In the ant *Myrmecina nipponica*, growth of the compound eye, forewing, and hindwing discs diverge in growth and final size between queens, workers, and a novel reproductive caste called ergatoid queens showing that disc growth is modular (Miyazaki et al., 2010). This demonstrates that in ant evolution, modifications to the growth and coordination relationships between discs and larval ontogeny generate the developmental differentiation of castes. These ancestral mechanisms of modularity and plasticity could have facilitated the acquisition of a novel regulatory role by the soldier rudimentary forewing discs in *P. hyatti*. Furthermore, the growth of rudimentary wing discs varies across other species with complex worker caste systems, in which minor worker larvae develop small wing pads and soldiers develop larger rudimentary wing discs and show inter-specific differences across species that lack soldiers yet develop larval wing rudiments (Abouheif & Wray, 2002; Bowsher et al., 2007; Shbailat & Abouheif, 2013; Rajakumar et al., 2018). Therefore, the diversity of morphology and elaboration of rudimentary wing discs across ants presents a useful system to investigate the origin, growth, and repurposing of rudimentary organs during development and evolution.

Collectively, this study shows how alterations to the growth trajectories of imaginal discs, including a rudimentary organ, by environmental cues can generate discrete and highly adaptive morphologies in ants. We show that the growth of the soldier rudimentary forewing discs is coordinated with the other discs and may have differential regulatory functions over the growth of the leg and eye-antennal discs and may regulate eye-antennal disc morphogenesis to generate disproportionate head-to-body scaling in the soldier subcaste in *P. hyatti*. By characterizing and describing the developmental processes that underlie these morphologies, we can begin to elucidate the processes through which worker subcastes differentiate and how disproportionate head-to-body allometry is generated during development. This highlights the emerging properties of this system for the study of rudimentary wing discs and their functional role in the development and evolution of worker caste systems across ants. More generally, future studies should examine the developmental underpinnings of the elaboration and function of rudimentary organs may reveal that they play a role in the regulation of novel phenotypes during development and evolution. Furthermore, future studies should also capitalize on nano-CT as a powerful technique to reveal the larval origin of many key adult features in ants, such as eyes, antennae, head capsule, mandibles, thorax, legs, and genitals, that are often elaborated and differentially expressed between queen and worker castes.

## Acknowledgments

We dedicate this manuscript to our collaborator Dr. Tomonari Kaji, a phenomenal researcher who passed away in May 2019 prior to the completion of this work. We also thank Bob Johnson for help collecting ants, Dr. Roberto Keller, and the Abouheif Lab for comments on this manuscript. This research was performed using the infrastructure of the Integrated Quantitative Biology Initiative (IQBI), Canadian Foundation of Innovation project 33122. We thank the McGill Cell Imaging and Analysis Network (CIAN) and the IQBI for equipment support and the Advanced BioImaging Facility (ABIF) for training. This work was funded by a National Sciences and Engineering Research Council of Canada (NSERC) Discovery Grant to E.A. and an NSERC Canada Graduate Scholarship (Master’s) and Fonds de Recherches de Quebec – Nature et Technologies (FRQNT) Scholarship to S.K.

## Notes

### Competing Interest Statement

The authors have declared no competing interest.

